# A gavage-fomite based method to generate mouse models with natural microbiota

**DOI:** 10.64898/2026.06.15.732393

**Authors:** Geongoo Han, Mohammad H. Hasan, Olumide Adesioye, Jessica Pacia, Julian C. Ramprashad, Gregorio Valdez, Shipra Vaishnava, Lalit K. Beura

## Abstract

Laboratory-raised specific pathogen-free (SPF) mice have been indispensable for fundamental immunology research, yet their reliability in predicting human clinical outcomes has been questionable. A major factor contributing to this disconnect is the sanitized housing environment, which deprives laboratory mice of physiological microbial exposure critical for immune maturation. Various approaches have been developed to introduce microbes to SPF mice, aiming to mimic human-like microbial experiences and engender adult human-like immune traits. However, some of these methods, specifically the pet store mice cohousing approach suffer from significant variability in pathogen exposure driven by the uncontrolled nature of microbial exchange and is associated with heightened mortality. Here we present an alternative gavage-fomite (GaF) method that exposes mice to a similarly diverse array of pathogens and commensal as the pet store cohousing method while limiting mortality. GaF-treated mice exhibited consistent gut microbial composition, robust immune maturation characterized by mucosal T-cell distribution, elevated serum inflammatory cytokines, and a splenic immune transcriptional signature closely aligned with that of adult humans. Furthermore, these mice demonstrated enhanced protection against a virulent bacterial challenge. The simplicity, and effectiveness of the GaF method for generating ‘mice with natural microbiota,’ may support broader use of these models in basic and translational immunological research across institutions.

## Introduction

Specific pathogen-free (SPF) laboratory mice have been the cornerstone of biomedical research and have provided crucial insights into fundamental biological processes since the last century. But their value in faithfully predicting human responses is questionable. SPF mice are housed in controlled environments, shielded from unintentional pathogen exposure and do not have the complex microbial experiences that shape human immune systems. Humans interact with a diverse array of pathogens, commensals, and environmental factors, creating immune responses that ultra-clean laboratory mice lack. The controlled nature of SPF models makes them valuable for isolating specific variables, but this comes at the cost of ignoring the broader biological complexities humans experience. These limitations underscore the need for alternative models and more integrated approaches that better replicate the multifaceted human experience.

In response to this need for better animal models, several groups have developed “mouse with natural microbiota” (MNM, a broader term suggested recently to encompass various so called “dirty mice” models) that intentionally expose laboratory mice to an array of microbes through various modifications in animal husbandry practices (1). The overall goal of this approach has been to generate mice with comparable microbial experience as adult humans and consequently with a matured/developed immune system. In the rewilding approach, laboratory SPF mice were placed in outdoor enclosures or semi-natural habitats, exposing them to environmental microbes over time (2, 3). Mice in this strategy must forage, interact with other animals, and survive without consistent human interventions. This allows for gradual colonization with a natural microbiota while maintaining some control over environmental variables. A similar approach termed feralization, housed SPF mice in barn enclosures along with wild-caught mice leading to acquisition of diverse microbial species (4, 5). More controlled approaches including the wildR and wildling mice involved either directly inoculating timed pregnant germ-free mice with wild mice microbiome or implanting SPF embryos in wild dams respectively (6, 7). All of these approaches generated mice with a rich microbiome and/or pathogenic exposure and mature immune phenotype. However, these protocols require access to specific infrastructure as well as specific technological expertise that all laboratories might not readily possess.

Several other approaches have also been developed that can be easily adapted across laboratories without significant technological challenges including the sequential exposure model and the pet store cohousing model. In the sequential exposure model, the investigator delivers a defined cocktail of pathogens to the SPF mice sequentially mimicking the early life pathogenic experience of humans (8, 9). In the pet store cohousing model, SPF mice are cohoused with pet store mice for a period of at least 60 days to allow transmission of pathogens as well as commensals from the pet store partner to the cohoused SPF mice (10–13). While the former approach can be more readily adapted because of limited biosafety requirements, the cohousing approach generates mice with broader infectious experience and a robust immune phenotype matching that of adult humans. However, cohousing with pet store mice generates mice in which the microbial exposure history can vary widely from cage to cage and even among animals in the same cage. Although this differential microbial experience could capture the spectrum of microbial heterogeneity humans encounter, it can also have differential impact on immune system that is hard to control. Recent work comparing several MNM models showed that different approaches can have strikingly distinct response to vaccination (14). Such variability can complicate interpretations and might hinder robust validation of studies across laboratories.

Here we have established a modified protocol for generating MNM that builds upon the pet store cohousing model but reduces the associated variability in pathogenic and commensal exposure. Importantly, this gavage/fomite (GaF) method exhibited similar immune maturation traits as described for the cohousing method but without the associated mortality.

## Results

### The gavage/fomite (GaF) method introduces broad pathogenic experience while limiting mortality

While cohousing mice in confined environments (cages) has been proved effective in transferring various microbial entities from ‘dirty’ pet store/wild mice to ‘clean’ laboratory mice, aforementioned limitations including variability in pathogenic load and mortalities upon exposure led us to investigate alternative approaches to deliver microbial experience. The primary route of microbial exposure in cohousing settings is thought to be through fecal-oral route mediated by coprophagy and grooming. We reasoned that the exposure to the cecal content of pet store mice along with the dirty bedding could substitute for the co-habitation-mediated microbial exposure. Briefly, we collected cecal contents of pet store mice (pooled from >30 donors) and administered orally to SPF C57BL/6 mice (see methods for details). The fomites, dirty beddings generated from the pet store mice cages, were also transferred to the gavaged mice cages twice per week throughout the experimental period (Fig. 1A). The fomite donors were replaced approximately every 5 weeks. We tested the ability of the gavage/fomite (GaF) method to successfully transfer common murine pathogens and compare it with cohoused animals by serological analysis (Fig. 1B). We also inquired if gavaging alone (Gav) will lead to comparable microbial experience as seen in GaF and cohoused (CoH) mice, as it would be a simpler method. Among the single-stranded RNA viruses tested, all MNM models showed equivalent seropositivity. However, only the GaF mice were seropositive for single-stranded DNA viruses including Minute virus of mice, Mouse Parvoviruses (type-1, 2, NS1), which are observed in pet store mice. On the other hand, a significant fraction of cohoused mice acquired the pathogenic bacterium, *Mycoplasma pulmonis*, from pet store mice while GaF and gavage-only groups remained seronegative. Among the murine pinworms, *Syphacia obvelata,* was detected in all MNM models, whereas *Aspiculuris tetraptera* was only detected in GaF group. Cohoused and GaF mice were positive for fur mites but gavage-only mice did not harbor fur mites. Our data suggest that the GaF method introduces a broad range of microbes including viruses, worms, and mites to laboratory mice as it was previously observed in pet store mice (10). The results also imply that fomite treatment is essential to expand the range of microbial exposure, especially non-intestinal microbes.

**Figure 1.**
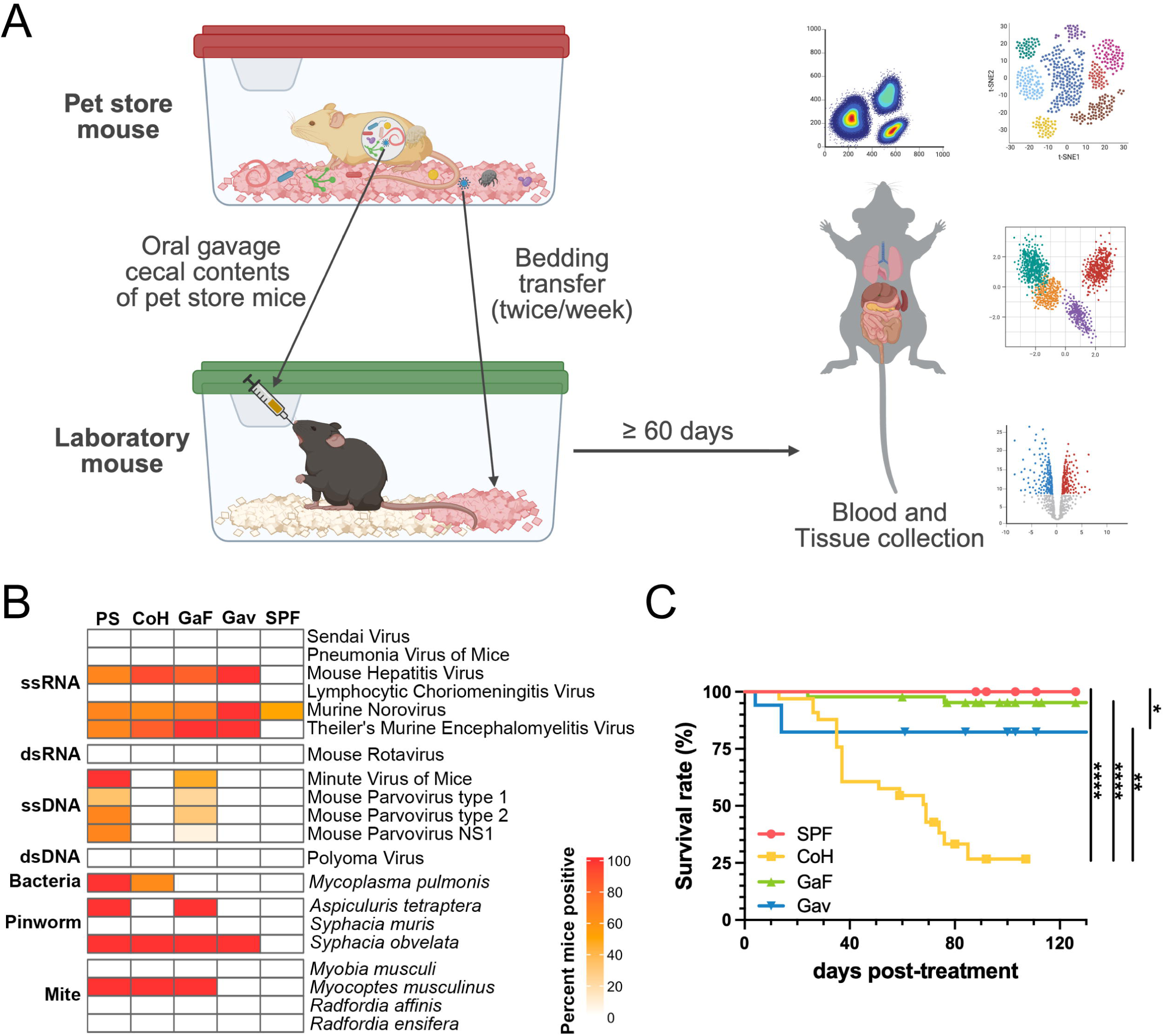
A comparison of survival rates and microbial exposure history across different MNM models. (A) Scheme of the Gavage/Fomite (GaF) method. Created with BioRender.com. (B) Heatmap showing distinct microbial experience among pet store mice (PS), cohoused (CoH), GaF, gavage-only (Gav), in comparison to specific pathogen-free (SPF) mice as determined by Charles River MFIA-based serology test and PRIA screening. (C) The survival rate of mice after treatment with various exposure regimens. Combined data from 3-5 independent experiments (CoH, *n* = 33; GaF, *n* = 45; Gav, *n* = 17). SPF mice were used as a control (*n* = 30). Significances of survival were determined by the log-rank (Mantel-Cox) test. *p < 0.05, **p < 0.01, and ****p < 0.0001.

Next, we compared the survival rates of various MNM models. The GaF group exhibited significantly higher survival rates (> 95%) compared with cohoused mice, over a 120-day observation period. Cohouse mice showed poor survival rates and more than half of them died within 40 days post-cohousing. Surprisingly, ∼18% of gavage-only mice also died within two weeks (Fig. 1C). Altogether, this suggests GaF technique is a robust approach to introduce diverse microbial experience without associated lethality seen in cohousing approach.

### Changes in the intestinal bacterial microbiome are more uniform in the GaF method than in cohousing

Gut microbiome has been recognized as a critical driver of immune fitness in various MNM models (2–13), but also could be a reason for variability. This variation can be pronounced in the cohousing approach as the microbiome composition of cohoused laboratory mice is largely influenced by the partner pet store mouse in the same cage. Herein, we decided to use petstore mice pooled gut content as the microbiome source material to be transplanted in SPF recipients. To test whether the GaF method can improve the uniformity in microbiome composition, we performed 16S rRNA gene sequencing and compared the intestinal bacterial microbiome among different MNM models.

The bacterial diversity within samples was comparable as checked by Shannon diversity, with no significant differences observed across the various MNM models and SPF mice (Fig. 2A). To assess the uniformity of the bacterial population, we evaluated within-group variability based on weighted UniFrac distances. Pet store and cohoused mice showed substantial variability in bacterial composition between individual samples, while the GaF and gavage-only groups exhibited lower within-group variability which was significantly lower than that observed in pet store or cohoused mice (Fig. 2B). In the principal coordinates analysis (PCoA) plot, SPF, GaF, and gavage-only samples clustered tightly, whereas pet store and cohoused samples were more variable (Fig. 2C). Next, we interrogated the relationship among the various dirty mice groups across separate cages. Interestingly, cohoused mice samples were separated by their cages and were clustered together with their other cage mates in the PCoA plot (Fig. 2D), whereas GaF mice did not show such segregation across cages. This suggests that gut bacterial composition in cohoused mice is influenced by the pet store partner mouse, leading to increased variability among cohoused mice.

**Figure 2.**
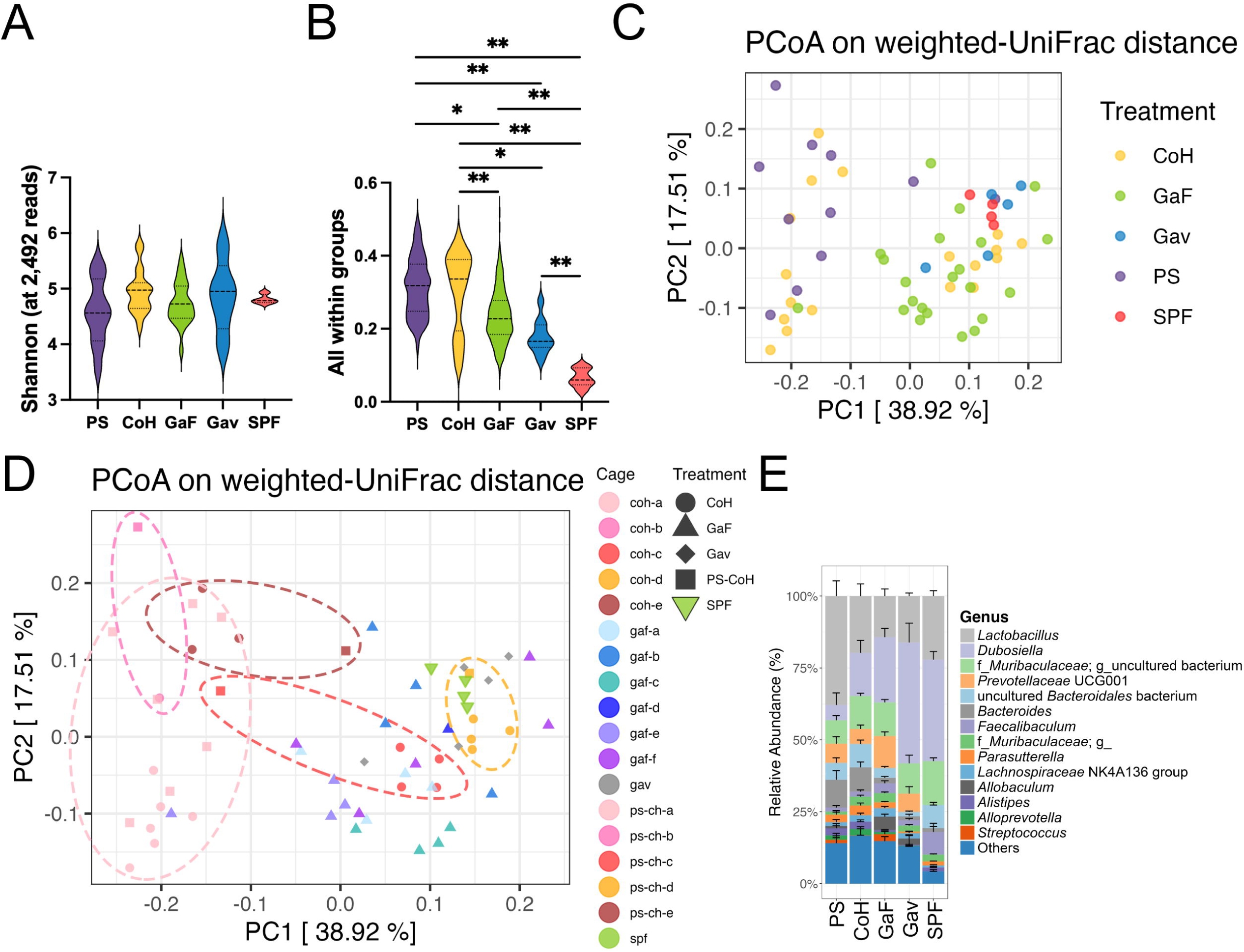
The bacterial community structure in the gut of different MNM models. Animals were treated with indicated microbial exposure methods, and feces were collected >60 days post-treatment to analyze bacterial diversity. (A) Alpha diversity based on the Shannon index. (B) Sample variability within the group based on weighted UniFrac distance. Significance was determined by PERMDISP. (C and D) PCoA plot based on weighted UniFrac distance. (C) Each mouse model is distinguished by color. (D) Cage-specific beta diversity. Each color represents different cages and each treatment group is distinguished by shape. (E) The relative abundance of the gut microbiome at the genus level. Bars indicate mean ± SEM. *p < 0.05 and **p < 0.01. PS: pet store mice, CoH: cohoused laboratory mice with pet store mice, GaF: gavage/fomite-treated mice, Gav: gavage-only mice, SPF: specific pathogen-free mice.

We next assessed the stability of the altered bacterial composition in GaF and gavage-only groups and found them to be maintained for at least 3 months (Fig. S1A). Interestingly, this alteration process was rapid and was already established 2 weeks post-treatment and changed minimally thereafter. We also compared the taxonomic composition of different mouse models at the genus level (Fig. 2E). Cohoused and GaF mice exhibited a similar genus composition to that of pet store mice, whereas notable differences were observed in gavage-only mice. Particularly, the relative abundance of *Bacteroides* was much lower in SPF and gavage-only mice, and *Dubosiella* was the most dominant genus in these mice, accounting for about 35% and 40% of total sequencing reads, respectively. Although GaF mice also exhibited a lower *Bacteroides* abundance, the overall genus composition was not much different from that of pet store mice or cohoused mice. Overall, the bacterial microbiome analysis suggested significant cage-specific variations among cohoused mice related to pet store partners and the gavage-only group showed interesting genus-level differences compared to other treatment groups.

### Immune phenotyping shows elevated antigen-experienced T cells and cytokines in GaF mice

A common aspect of increased microbial exposure across different MNM models is the accumulation of effector-biased immune cells in the systemic circulation (2, 4, 5, 9–13). Majority of MNM models have shown alteration in T lymphocytes and granulocytes (1–5, 7, 9–13). Here, we performed flow cytometry-based immunophenotyping of peripheral blood leukocytes (PBL) to characterize this effector differentiation. The t-distributed stochastic neighbor embedding (t-SNE) analysis was used to visualize overall immune cell compositions among the different approaches. We noticed substantial increase in antigen-experienced CD8^+^ T cells (CD44^hi^ CD8^+^ T cells) as well as KLRG1^+^ CD8^+^ T cells as early as 30 days-post treatment in all the MNM groups compared to the SPF mice (Fig. 3A). Frequency and numbers of these effector-biased CD8^+^ T cells also remain elevated when checked ≥ 60 days post-treatment (Fig. 3B and C). We further validated that a separate pool of pet store mouse cecal content (collected after an interval of >2 years) can similarly elevate the frequency of antigen-experienced CD8^+^ T cells, establishing the robustness of the method (Fig. 3D). Interestingly, the tSNE plots at day 60 (Fig. 3E top row) were markedly different than day 30 (Fig. 3A) and the day 60 CoH plot showed a highly abundant neutrophil and Ly6C^hi^ inflammatory monocytes population compared to all other groups. Our serological survey data suggest that this neutrophilia is driven by *M. pulmonis* infection, as cages of cohoused mice that were seronegative for *M. pulmonis* did not show elevated neutrophil level (Fig. 3E, bottom row). It is important to point out that both the cohoused mouse in Fig.3A as well as in Fig.3E (top row) were seropositive for *M. pulmonis,* but the increase in neutrophils was only evident in the >day 60 group suggesting the neutrophilic response might arise in response to chronic colonization. Interestingly, *M. pulmonis* seropositivity was above 64% among cohoused mice in our facility and we used *M. pulmonis* positive cohoused mice for all subsequent studies (Fig. 1B). The increased number of activated CD8^+^ T cell populations was sustained for at least 6 months (Fig. S1B and C).

**Figure 3.**
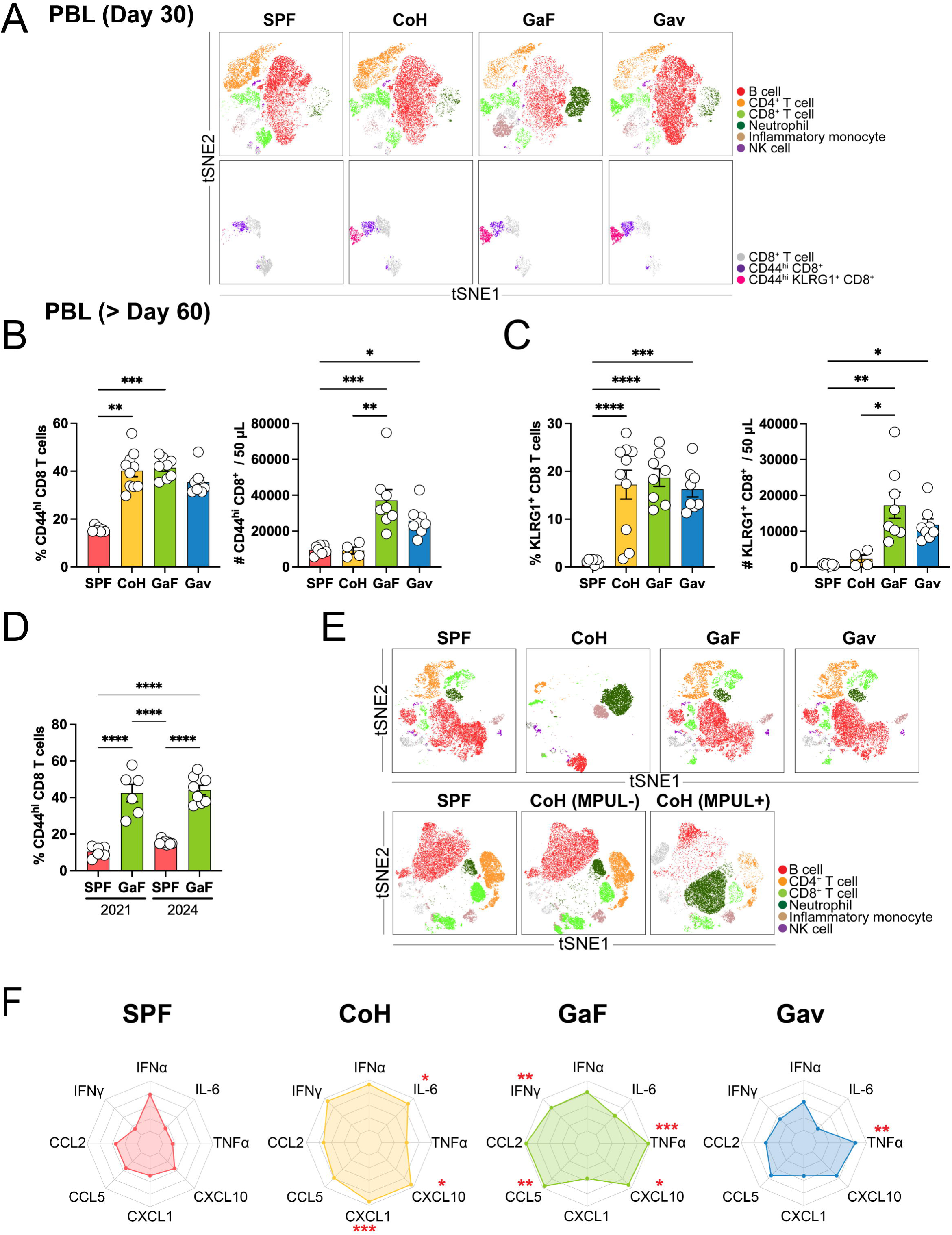
The immune cell composition of different natural microbiota-based mouse models in the blood. (A-D) Flow cytometry was used to characterize peripheral blood immune cells. (A) Representative t-SNE plot of the immune cells (gated on CD45^+^) and CD8^+^ T cells showing KLRG1 and CD44 expression (gated on CD45^+^ CD11b^-^ B220^-^ CD4^-^ CD8^+^) in the peripheral blood of SPF mice and different MNM models at 30 days post-treatment. Enumeration of CD44^hi^ CD8^+^ T cells (B) and KLRG1^+^ CD8^+^ T cells (C) frequencies and counts in blood >60 days post-treatment. (D) Frequency of CD44^hi^ CD8^+^ T cells in GaF mice blood prepared from two separate source of pet store mouse cecal content collected in year 2021 and 2024. (E) Representative t-SNE plot of CD45^+^ immune cells in the peripheral blood of SPF mice and various MNM models (top) and *Mycoplasma pulmonis* seronegative (MPUL-) and seropositive (MPUL+) cohoused mice (bottom) >60 days post-treatment. (F) Cytokine levels in the serum of SPF, CoH, GaF, and Gav mice. Combined data from 2 independent experiments. The log10 (average value) was used for the radar chart. Bars indicate mean ± SEM. Significances were determined by the Kruskal-Wallis test with Dunn’s multiple comparisons test or the one-way ANOVA test with Tukey’s multiple comparison test. In Fig. 3F, multiple comparison tests were performed between SPF and each natural microbiota-based mouse model to find significant differences. *p < 0.05, **p < 0.01, ***p < 0.001, and ****p < 0.0001.

We also noticed significant reduction in the B cell population in the CoH mouse PBL compared to all the other mouse groups (Fig. 3E). To understand, how this change might alter different subsets of blood-borne B cells, we performed flow cytometry based enumeration of naïve and memory B cells in PBL. We found a significant reduction in the frequency of naïve and unswitched B cells in CoH group compared to all the other mouse groups, although this difference was numerically insignificant from SPF group (Fig. S2A). On the other hand, both frequency and number of switched memory B cells was higher in all the MNM mice compared to SPF mice (Fig. S2B).

We also sought to test whether the GaF treatment can be applied to male mice, because a key limitation of the cohousing method is it cannot be applied to male mice as male cohousing beyond the weaning age often results in aggressive behaviors among cage mates. The GaF-treated male mice also showed similarly increased levels of differentiated T cell populations as that of female mice (CD44^hi^ and KLRG1^+^ CD8^+^ T cells) (Fig. S3A – C).

Fomite treatment alone has been shown to be a less robust method in mediating immune maturation (15). We further tested this assertion in our facility. We established a separate cohort of mice receiving only fomites twice weekly (Fom) and checked the frequency of antigen-experienced cells in the spleen. Although fomite treatment alone increased the population of CD44^hi^ CD8^+^ T cells, the level of increase was much lower than that observed with other methods (Fig. S4A and B). This suggests that additional microbial exposure beyond fomite is necessary to achieve more robust activated CD8^+^ T cell phenotypes.

Another aspect of immune maturation in MNM mice is the elevated presence of certain cytokines than their SPF counterpart. Cohousing increased levels of several inflammatory cytokines including CXCL10 and neutrophil chemoattractants like CXCL1 and IL-6 compared to SPF mice as has been described before (13). GaF mice also showed significantly higher levels of inflammatory cytokines than SPF mice, including IFN-γ, CCL5, CXCL10, and TNF-α. In contrast, the gavage-only mice exhibited an increase in TNF-α level alone in peripheral blood (Fig. 3F and Fig. S5). Collectively, the GaF method generated and sustained increased frequency of effector-like CD8^+^ T cells, memory B cells similar to cohoused mice and also showed increased levels of several proinflammatory cytokines and chemokines. Presence of *M. pulmonis* in cohoused mice was associated with neutrophilia, which was absent in GaF mice.

### Mucosal barrier organs show evidence of immune infiltration in GaF mice

A cardinal feature of immune maturation is the generation of activated innate as well as adaptive immune cells that then get distributed in various lymphoid and nonlymphoid tissues across the body. Here, we examined whether the GaF method is capable of depositing immune cells across a number of mucosal tissues. Using a spectral flow cytometry approach that is capable of distinguishing at least 19 different populations of immune cells (an example gating scheme is shown in Fig. S6), we examined the immune composition of the small intestine epithelium. Intestinal epithelial leukocyte population was dominated by CD8^+^ and CD4^+^ T cells in mice that were exposed to pet store mice-derived microbes compared to SPF control as shown in the t-SNE plots (Fig. 4A). We also enumerated the number of these different populations and depicted the abundance of different immune populations using heatmaps. GaF, cohoused, and gavage-only mice clustered together, segregated from SPF mice based on the number of these various immune cells (Fig. 4B). However, among the various microbe-treated groups, the number of resident memory-like CD8^+^ T cells was highest in GaF and cohoused mice than the gavage-only group. Similar to the gut, there was also increased density of differentiated effector and resident memory-like T cells in the lung parenchyma of microbe-exposed animals (Fig. 4C and D). Interestingly, the cohoused mice showed elevated frequency of macrophages and neutrophils in the lung compared to other groups, similar to the rise in the percentage of these cells in systemic circulation (Fig. 3E and 4C).

**Figure 4.**
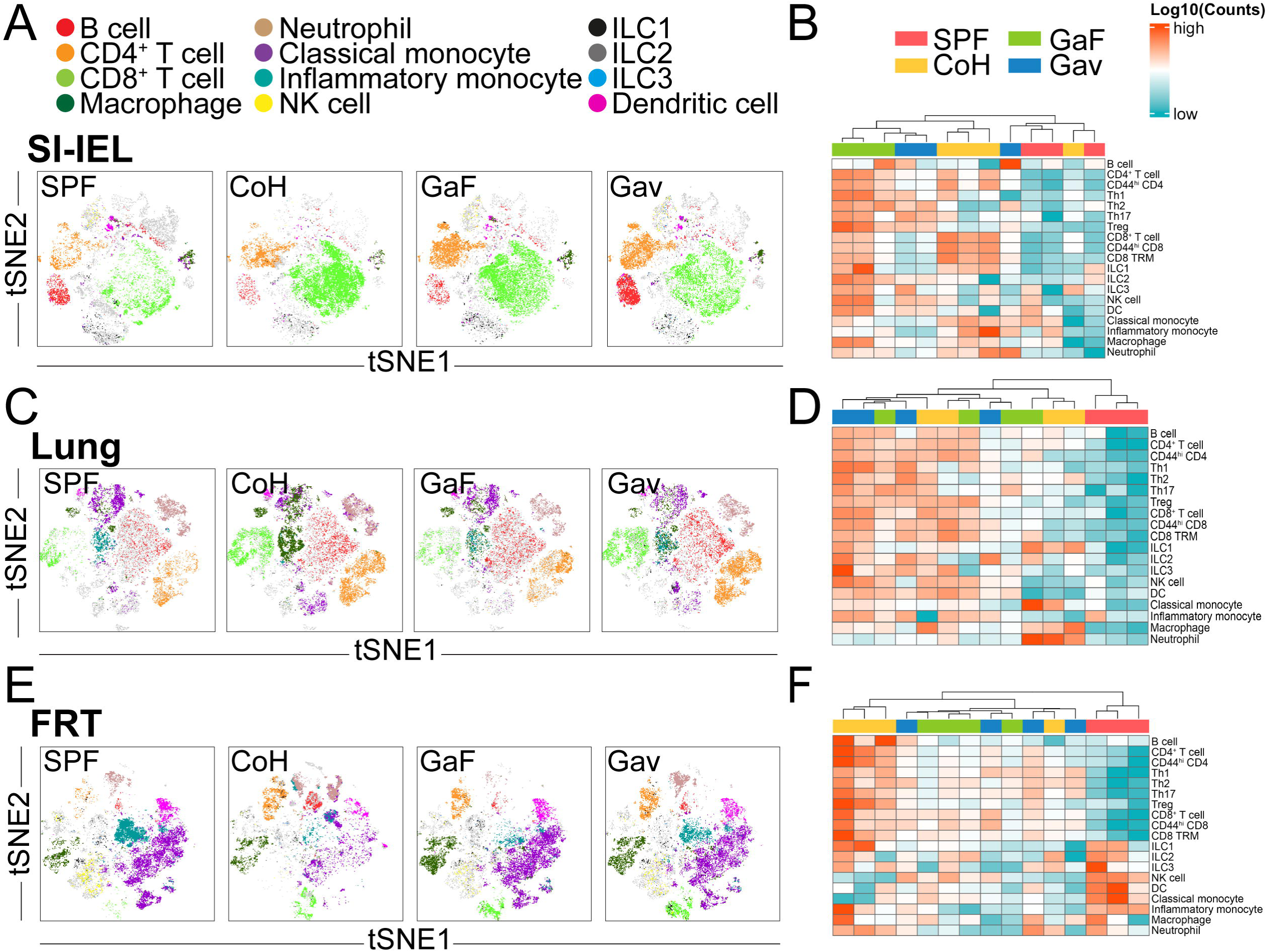
The immune cell composition of mucosal tissues across different MNM models. Animals were treated with different microbial regimens to generate mice with natural microbiota as described. Tissues were harvested >60 days post-treatment and flow cytometry was used to characterize immune cells. t-SNE plot showing relative abundance and distance among various immune cells in the small intestine epithelium (SI-IEL; A), lung parenchyma (C), and female reproductive tract (FRT; E). Indicated cell types were enumerated, and number of cells were log_10_ transformed and normalized per row and is depicted in the heatmap (B - SI-IEL; D - lung parenchyma; F - FRT). Heatmap rows indicate cell types, and columns represent individual mice.

We tested the immune composition of the female reproductive tract (FRT) and noticed a significant rise in the frequency of CD4^+^ and CD8^+^ T cells in GaF, cohoused, and gavage-only mice compared to SPF mice that barely had any T cells (Fig. 4E). We noticed that among the various microbial exposure regimens, the cohoused mice FRT contained highest numbers of all CD4^+^ T cell lineages compared to GaF and gavage-only mice (Fig. 4E and F). This might be indicative of specific microbial signals available in the cohoused setting that is limited in other methods. Altogether these immunophenotyping analyses showed increased generation of effector immune cells upon microbial exposure that were broadly distributed across the mouse body.

### Transcriptional changes in GaF showed enrichment of type-1 IFN signature

Our results indicate that the GaF method alters immune cell populations similar to cohoused mice. Because the cohousing approach has been known to drive an adult human-like immune signature in mice (10), we performed bulk RNA-seq of splenocytes in GaF group and compared it with previously published gene expression profiles of cohoused mice (GSE78979)(10) and human PBMCs (GSE27272)(16). The gene set enrichment analysis (GSEA) revealed the gene expression patterns of the GaF mice in the spleen enriched for both adult human PBMC (NES = 1.3689, Padj = 0.0075) and cohoused mice (NES = 1.5381, Padj = 0.0006) transcriptional signatures (Fig. 5A and S7A). Interestingly, when we tested if mice that were only gavaged with pet store microbiome without any fomite treatment can have similar cohoused/adult human immune gene enrichment, the transcriptome of gavage-only group failed to overlap significantly with that of cohoused (NES = 1.0658, Padj = 0.6722) and adult human PBMC (NES = 0.7295, Padj = 0.9876) (Fig. 5B and S7B). We also performed the gene ontology (GO) enrichment analysis to characterize the transcriptome following the GaF treatment. Notably, several GO terms related to type I interferon (IFN) responses were enriched in the GaF mice (Fig. 5C) as has been seen in past cohoused studies (10). We observed that several type I IFN-related genes such as *Arg1*, *Igtp*, *Ifit1*, *Irf7*, *Oas1a*, *Oas2*, *Ifitm3*, and *Pycard* were significantly higher in the GaF mice than SPF mice, and those genes were also highly expressed in the adult PBMC than in the neonatal PBMC (Fig. 5D). Taken together, these findings indicate that the GaF method better approximates the transcriptional makeup of adult humans similar to cohoused mice dominated by a type I IFN response. Whereas similar transcriptional alterations in gavage-only mice are insignificant.

**Figure 5.**
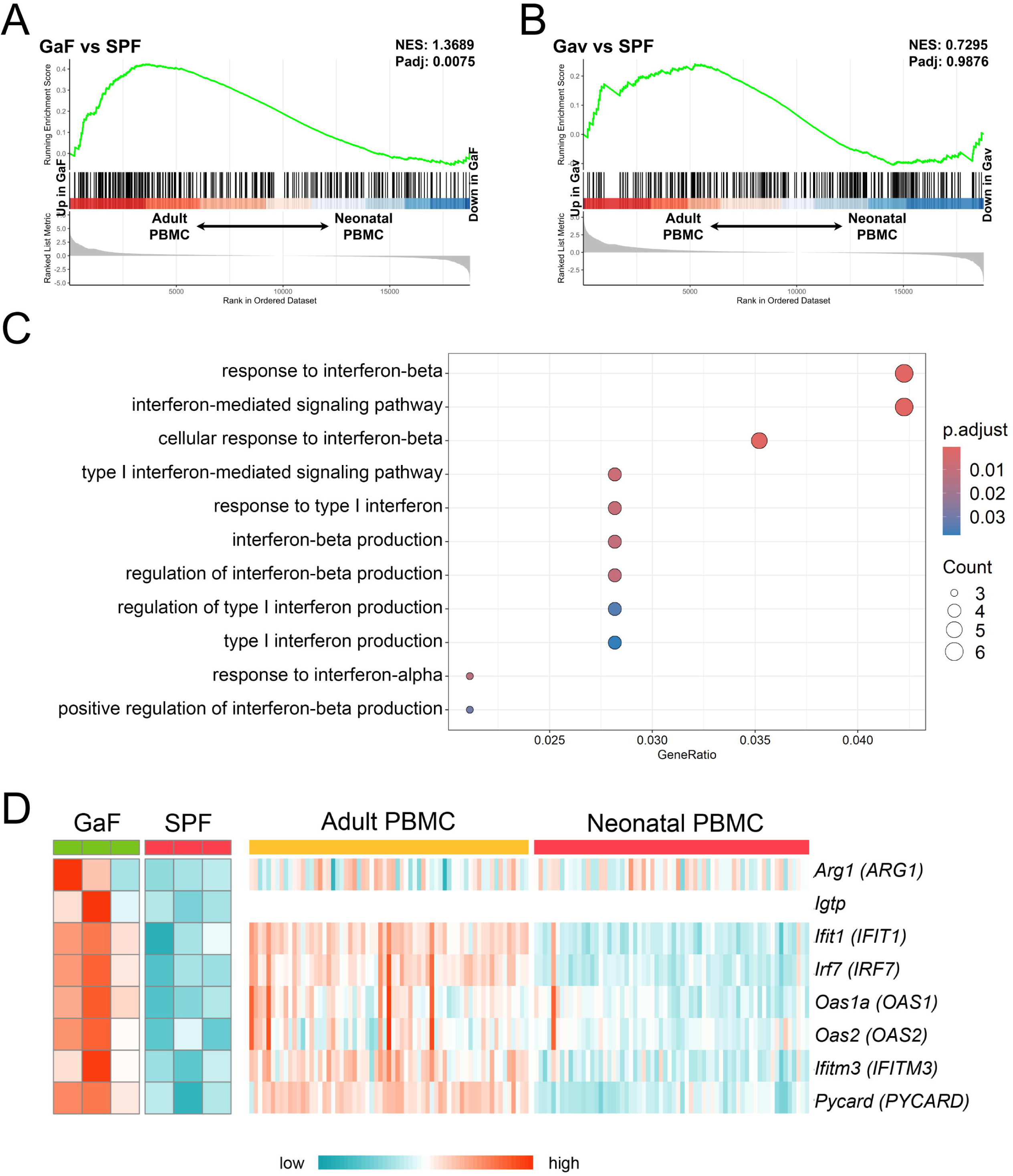
Transcriptional overlap between GaF mice and adult humans. Transcriptional profiles of GaF and gavage-only mice were generated by bulk RNA-sequencing of splenocytes and were compared with existing adult and neonatal human PBMC datasets. (A) GSEA plots of GaF vs SPF and (B) Gavage-only (Gav) vs SPF showing relative enrichment of differentially expressed genes (DEGs) between adult and neonates. The signature was derived from the top 400 DEGs (data obtained from GSE27272). (C) Gene Ontology (GO) term enrichment analysis. Significant type I interferon-related GO terms were visualized in the dot plot. (D) Heatmap of the type I interferon-related genes based on the GO analysis above. Values are log10 (normalized counts).

### GaF mice with matured immune repertoire are protected against a virulent *Listeria* challenge

Next, we assessed the functional capability of the matured immune system of GaF mice by using a virulent *Listeria monocytogenes* systemic infection that is fatal to SPF mice (Fig. 6A). *L. monocytogenes*-challenged SPF mice experienced heightened mortality, whereas the GaF and pet store mice showed complete survival (Fig. 6B). We also measured daily body weight of these L. monocytogenes-challenged mice and found that the GaF mice showed a small and transient weight loss around day 1-2 post-challenge, but their weights were quickly stabilized (Fig. 6C). However, the SPF group continued to lose weight (Fig. 6C). The pet store mice didn’t experience any weight loss. In a separate cohort of animals, we also estimated *L. monocytogenes* burden in the liver. The bacterial load was also significantly lower in the GaF mice than in the SPF mice, and it was comparable to that of pet store mice (Fig. 6C).

**Figure 6.**
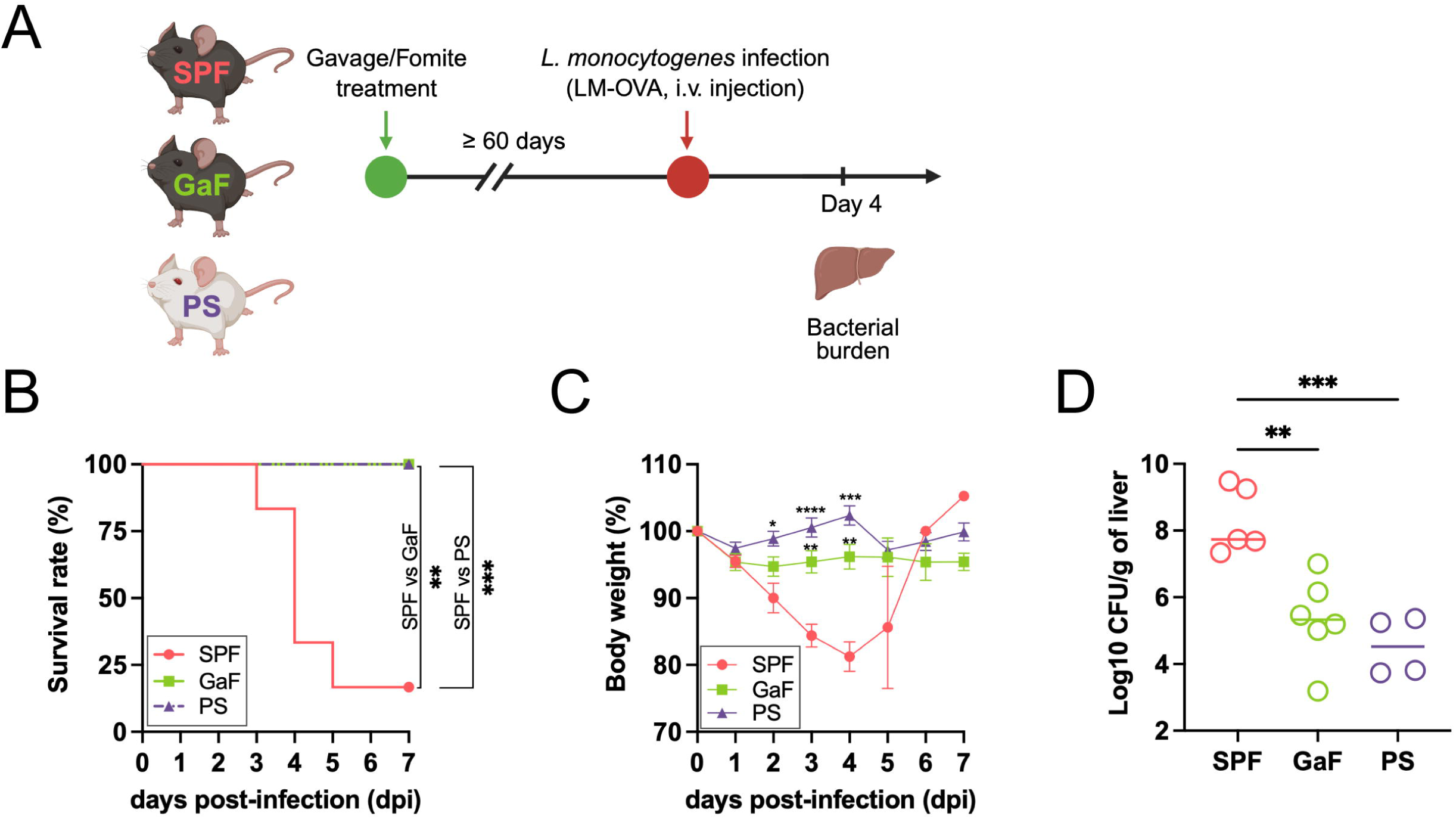
The gavage/fomite-treated mice are significantly protected against *L. monocytogenes* infection. (A) Scheme of the infection. SPF and pet store mice (PS) were used as controls. Created with BioRender.com. (B) Survival rate after *L. monocytogenes* infection. Significances of survival were determined by the log-rank (Mantel-Cox) test. (C) Body weight changes after *L. monocytogenes* infection. Statistical significance was determined by a mixed-effects analysis with Tukey’s multiple comparison test. Asterisks indicate significant differences between SPF mice and either GaF or pet store mice. (D) Bacterial load in the liver on day 4 after the challenge. Bacterial load was normalized by the weight of the liver. Significances were determined by the one-way ANOVA test with Tukey’s multiple comparison test. *p < 0.05, **p < 0.01, ***p < 0.001, and ****p < 0.0001.

## Discussion

Among vertebrates, the laboratory mouse (*Mus musculus*) has long been a model of choice for many human diseases of infectious, hereditary, and metabolic origins (17, 18). Within the field of immunology and infectious diseases, the mouse has become a cornerstone of research largely because of the ease of genetic manipulation, their relative affordability, and the availability of associated reagents to conduct reductionist experiments. Despite the power of mouse-based immunological research, there have been many examples wherein the use of laboratory mouse models resulted in misdirection in clinical research associated with significant economic costs and failure to positively impact human suffering. These failures have often been explained by poor experimental design, imperfect recapitulation of disease pathogenesis, and differences in underlying biological networks that control the disease process. As such there is a growing call to refocus experimental efforts on humans. While the rapid growth of the systems biology field and the accessibility of associated multi-omics technologies have made it easier to conduct experiments in humans, there are many experiments that can only be performed in a model organism because of technological and ethical reasons. Accordingly, there is an emergent need of developing mouse models that better recapitulate the human condition.

Humans just like any other free-living species are routinely exposed to pathogenic and commensal microbes (part of our “metagenome”) even before birth, which along with intentional exposures like immunizations, contribute to maturation of our immune system. In contrast, laboratory mice are actively prevented from being exposed to natural mouse microbes which greatly impairs the maturation of their immune system (10). Cohousing SPF mice with pet store mice transferred the natural microbiome to SPF laboratory mice and brought their immune cell composition, phenotype, and immune transcriptome closer to those of humans (10). This along with additional models developed across a number of laboratories suggest a critical contribution of a normalized microbial exposure in shaping our immune compartments (1–13). Importantly, these changes significantly altered kinetics of diseases, immune responses as well as responses to vaccines and therapies aligning more closely with human responses (6, 7, 14, 19). However, a critical limitation of some MNM models is the inherent variability in microbial experience across the animals. While this variation is probably similar to human microbial exposure differences, the unknown nature of the variables will prevent drawing robust conclusions. This variability issue is most acute in the pet store cohousing and feralization approaches as the microbial burden and composition in the pet store and wild mice are uncontrolled. Another drawback of the pet store cohousing method is the high mortality of the laboratory mice which can range from 10% - 100%. We have reported an average of 25% in an earlier study (10), but in the current study, this value was closer to 70%. Here, we described a normalized microbial exposure technique-GaF method that reduces microbial exposure variability by bulk sourcing of the gut microbiome from pet store mice along with constant microbial exposure via biweekly fomite treatment. GaF-treated mice also showed excellent survival (≥ 95%). The GaF-treated animals displayed robust immune maturation with differentiated effector immune cells distributed across peripheral tissues, exhibiting a type I IFN-dominated transcriptional signature similar to cohoused mice. The GaF mice also showed elevated inflammatory cytokine/chemokine levels and survived a virulent *L. monocytogenes* challenge.

The commensal bacteriome is a crucial component of our metagenome with significant influence on host physiology and immunity. Our 16S rRNA analysis showed marked cage-to-cage variability in bacterial composition in the cohoused treatment group. As expected, this difference was primarily driven by the pet store partner mouse. This differential microbial composition could also influence immune composition, as seen in the immunophenotyping studies. In comparison, when we measured bacterial homogeneity, we found that the GaF group had a more homogeneous bacterial composition compared to cohoused mice. Interestingly, the alteration in bacterial community structure introduced by the gavage process was fairly rapid (notable within 2 weeks) and remained stable for at least 3 months even though the mice were regularly exposed to new pet store mouse beddings drawn from different pet store mouse cages over time. This attests to the dominant nature of the non-SPF microbiome.

In an effort to check if delivery of pet store gut content alone without the fomite treatment can drive similar immune activation, we thoroughly characterize the gavage-only group. Interestingly gavage-only mice did show increased activated T cells in blood and mucosal tissues, but the systemic cytokine/chemokine analysis showed a TNF-α specific response in the absence of other inflammatory molecules like IFNγ, CXCL10 which were also present in GaF mice. We also noted that gavage-only mice did not show enrichment of adult human-like immune signature unlike cohoused and GaF groups. Interestingly, the serological survey showed that gavage-only group were seronegative for all the ssDNA viruses as well as mites and certain species of pinworms. Lack of this microbial exposure could be the reason behind incomplete immune maturation as well as the heightened mortality (≥ 18%) experienced by the gavage only group. The regular fomite treatment is an important substitute for routine microbial exposures we experience through the life. But fomite treatment alone did not induce robust immune maturation.

Although our GaF technique reduced variability in microbial exposure and improved survival, several limitations remain. The approach still relies on non-SPF source material that is incompletely defined in terms of exact microbial makeup. As such, this approach will still require specialized housing setup to prevent accidental spread of pathogens to normal SPF mouse housing. We were able to house these animals in a biosafety level-2 housing that is far away from regular animal facilities. But these might require significant institutional commitments. Identification of the ideal microbial composition that can mediate robust and reproducible immunological changes will streamline the process, reduce cost and enable wider usage. A microbiologically stable model like GaF mice will be an important starting point in curating such a list of microbes with well-characterized transmission and pathogenesis profiles to overcome this issue.

In summary, we have provided a tool to generate a mouse model with natural microbiota with broad pathogenic experience that can be adopted easily. Detailed characterization of this GaF model showed that the mice showed superior survival and equivalent immune maturation profile to that of the popular pet store cohousing model. We were able to reduce variation in pathogen and commensal exposure, which led to robust and sustained immune activation. Use of this model will enable a better understanding of the human immune system using laboratory mice with reliable and reproducible results across institutions.

## Materials and methods

### Mice

Specific pathogen-free (SPF) laboratory C57BL/6 mice were purchased from Jackson Laboratory and bred in the animal facility at Brown University. Pet store mice were purchased from local pet stores in the Rhode Island area and were housed in a specific investigator-managed animal housing space separate from routine laboratory animal housing space. For this study we have primarily focused on female mice, but male mice were used in experiments comparing immune maturation in male vs female animals. For the cohousing group, one female pet store mouse was cohoused with four female SPF mice in the same cage. For the fomite treatment, fomite cages were maintained with 5-6 pet store mice per cage, and dirty beddings (∼12 g) were transferred from pet store mice cages (maintained in our facility) to SPF mice cages (75 square inches) twice per week throughout the whole experimental period. For collection of the cecal contents, a group of >30 pet store mice were euthanized and the cecal contents were harvested immediately followed by resuspension in sterile PBS. The suspension was filtered (70 μm) and the filtrates were mixed with 50% glycerol (1:1) and were stored at −70 °C until oral gavage. 100 μL glycerol stocks were orally gavaged to SPF mice at the beginning of the experiment. All treated mice were euthanized after at least 60 days of treatments. Experiments were performed according to the protocols approved by the Institutional Animal Care and Use Committees of Brown University.

### Microbial exposure estimation

Virus and bacterial exposures were determined by Multiplexed Fluorometric ImmunoAssay (MFIA)-based serology test (Charles River Laboratories) on blood/serum. Pinworms and mites exposure were screened by PCR Rodent Infectious Agent (PRIA) test (Charles River Laboratories). Fecal samples and fur swabs were collected for PRIA tests. Samples were collected from cohoused, GaF, and gavage-only mice after at least 50 days of treatments.

### 16S rRNA gene sequencing and analysis

Total gDNA was extracted from fecal samples using a Quick-DNA Fecal/Soil Microbe Microprep Kit (Zymo Research) according to the manufacturer’s protocol. The bacterial V4/V5 region of the 16S rRNA gene was amplified using the Phusion High-Fidelity DNA polymerase (Thermo Scientific) with 518F/926R primer pair (518F: 5’-CCAGCAGCYGCGGTAAN-3’, 926R: 5’-CCGTCAATTCNTTTRAGT-3’), which were conjugated with overhang adapter sequences for Illumina MiSeq with following PCR program: 95 °C for 3 min, followed by 35 cycles of 95 °C for 30 sec, 58 °C for 30 sec, and 72 °C for 30 sec, and 72 °C for 10 min. DNA libraries were constructed and sequenced on the Illumina MiSeq platform (paired-end 300 bp) at Rhode Island Genomics and Sequencing Center.

QIIME2 pipeline was used for microbiome analysis (20). The raw sequencing reads were trimmed and denoised followed by amplicon sequence variants (ASVs) table construction using DADA2 (21). Taxonomy was assigned using the pre-trained Naïve Bayes classifier on the SILVA 132 database (22). Singletons and all features annotated as mitochondria or chloroplast sequences were removed from the ASV table and the bacterial abundances were expressed as a percentage of total 16S rRNA gene sequences. For alpha and beta diversity, the feature table was rarefied to 2,492 reads. Shannon was used as an alpha diversity index and principal coordinates analysis (PCoA) based on weighted or unweighted UniFrac distances was used for beta diversity. PCoA results were plotted using the R package *qiime2R* (https://github.com/jbisanz/qiime2R). PERMDISP was used to find significant differences in sample dispersion within the group.

### Immune cell isolation and phenotyping

Blood samples were collected by cheek puncture in heparin-containing polystyrene tubes to isolate the peripheral blood leukocytes. Red blood cells were lysed by treatment with ACK lysis buffer.

Before tissue harvests, mice were injected intravenously with fluorochrome-conjugated anti-CD45 to distinguish cells in the vasculature and the tissue parenchyma. Three minutes postinjection, mice were euthanized, and tissues were collected. Immune cells in the spleen, small intestinal epithelium (SI-IEL), female reproductive tract (FRT), and lung were isolated as described in Beura et al. (23). Briefly, for isolation of SI-IEL, Peyer’s patches were removed, and the SI was cut longitudinally and then laterally into small pieces. Pieces were incubated for 30 min with shaking at 37 °C with 0.154 mg/mL dithioerythritol (MilliporeSigma) in 10% HBSS/HEPES. The pieces were vortexed on high speed to dislodge IELs and the IELs were collected for further density gradient centrifugation. Lung and the FRT (containing uterine horns, cervix, and vaginal tissue) were removed and cut into small pieces, followed by treatment with 1 mg/ml of type IV (MilliporeSigma) collagenase in 5% RPMI-1640/ 2 mM MgCl_2_/ 2 mM CaCl_2_ (45 min for lung and 1 h for FRT at 37 °C, 250 rpm). All tissue pieces were washed several times before enzymatic digestion with 5% RPMI-1640 medium to remove excess of injected antibodies so as to prevent staining extravascular cells. IELs were purified on a 44%/67% Percoll gradient (800 *xg* at 23 °C for 20 min).

Isolated cells were stained with the corresponding cocktail of fluorescently labeled antibodies (CD4, CD8a, CD11b, CD11c, CD19, CD26, CD38, CD43, CD44, CD45, CD45.2, CD62L, CD64, CD69, CD90.2, B220, F4/80, GL7, IgD, IgM, KLRG1, Ly6C, Ly6G, MHC II, Nkp46, TCRβ). Cell viability was determined using Ghost Dye 780 (Tonbo/Cytek). After surface staining, cells were fixed and permeabilized using the Perm/Fix kit (Tonbo/BD) by following the manufacturer’s instructions. Intracellular staining with cocktail of fluorescently labeled antibodies (T-bet, Eomes, Foxp3, GATA3, Rorγt). The stained samples were acquired using the Aurora flow cytometer (Cytek). Flow cytometry data were gated and analyzed with FlowJo 10 software (BD). To visualize high-dimensional flow cytometry data, live CD45.2^+^ cells were sampled using the DownSample plugin and analyzed using the t-distributed stochastic neighbor embedding (t-SNE) method in Flowjo 10 software. The antibodies were purchased from Biolegend, Invitrogen, or BD Life Sciences.

### Antiviral cytokine assay

Blood samples were collected from the submandibular vein by cheek punch, followed by 1.5h incubation at room temperature and centrifugation at 2,000 *xg* for 15 min to separate the serum. The collected serum samples were then stored at −70 °C until further analysis. Anti-viral cytokines in the serum samples were quantified using LEGENDplex Mouse Anti-Virus Response Panel (BioLegend). R package *fmsb* was used to draw the radar chart.

### RNA-Seq

Small pieces of the spleen were stored in RNAlater (Invitrogen) and stored at −70 °C until RNA extraction. Total RNA was extracted using RNeasy Plus Micro Kit (Qiagen) according to the manufacturer’s protocol. RNA libraries were constructed from the extracted RNA samples and sequenced on the Illumina NovaSeq X Plus platform (paired-end 150 bp) at Novogene.

Adapter sequences and low-quality reads were trimmed from the raw sequence reads using Trimmomatic (24). Trimmed sequence reads were aligned to the mm10 mouse genome using the STAR aligner (25). Differentially expressed genes (DEGs) were distinguished if *Padj* < 0.05 in R package *DESeq2* (26). R package *clusterProfiler* (27) was used to perform gene set enrichment analysis (GSEA) and gene ontology (GO) enrichment analysis. For GSEA, we obtained the non-normalized human peripheral blood mononuclear cells (PBMCs) microarray dataset from the GEO database (GSE27272). This dataset includes samples from 65 adult women and 64 cord blood samples from their neonates. We also obtained the non-normalized mouse PBMCs microarray dataset (GSE78979). This dataset includes samples from 7 cohoused and 8 SPF mice. The human and mouse datasets were processed in *DESeq2*, and the top 400 upregulated genes in human adult or cohoused mice’s PBMCs were used as a signature dataset for GSEA. R package *Complex heatmap* (28) was used to draw the heatmap.

### L. monocytogenes challenge

Recombinant *L. monocytogenes* strain LM-OVA was cultured in brain heart infusion (BHI) media supplemented with 50 μg/mL streptomycin. The bacterial culture was grown until it reached an OD600 of approximately 0.1, then washed twice with sterile PBS. SPF, GaF, and pet store mice were infected intravenously with 1-3 × 10^5^ CFU (per mouse) LM-OVA.

The infected mice were euthanized 4 days post-infection, and the liver samples were collected. The livers were weighed and homogenized in 1 mL PBS containing 0.2% IGEPAL (CA-630) to lyse the cells. The homogenized liver samples were then serially diluted with sterile PBS and plated on the BHI agar plates supplemented with 50 mg/mL streptomycin. Following overnight incubation at 37 °C, colonies were counted to quantify the bacterial load in the liver.

## Statistical analysis

All statistical analysis and plotting were performed on R (https://www.R-project.org) and Prism (GraphPad). The normality of data was assessed by the Shapiro-Wilk test. If data followed a normal distribution, parametric tests (unpaired two-tailed t-test for two groups, One-way ANOVA with Tukey’s multiple comparisons test for more than two groups) were performed. If data did not follow a normal distribution, non-parametric tests (Mann-Whitney test for two groups, Kruskal-Wallis test with Dunn’s multiple comparisons test for more than two groups) were performed to find significant differences between groups. For the survival curve, the log-rank (Mantel-Cox) test was used to find significant differences between groups. *p < 0.05, **p < 0.01, ***p < 0.001, ****p<0.0001.

## Acknowledgments

Schematics were generated with Biorender.com. We acknowledge the Brown University flow cytometry core for help with the flow-based assays.

## Author contributions

Conceptualization: G.H., M.H.H., and L.K.B.; Methodology: G.H., M.H.H., and L.K.B.; Investigation: G.H., M.H.H., J.P., J.C.R., and L.K.B.; Funding acquisition: G.V. and L.K.B.; Resources: M.H.H., S.V., and L.K.B.; Writing – original draft: G.H. and L.K.B.; Writing – review & editing: G.H. and L.K.B.

## Data availability statement

Raw sequence reads from 16S rRNA gene sequencing have been deposited in the NCBI Sequence Read Archive (SRA) under accession number PRJNA1189529. Bulk RNA-seq data has been deposited in the NCBI GEO database under accession number GSE284503.

