## Supplementary information for "A gavage-fomite based method to generate mouse models with natural microbiota"

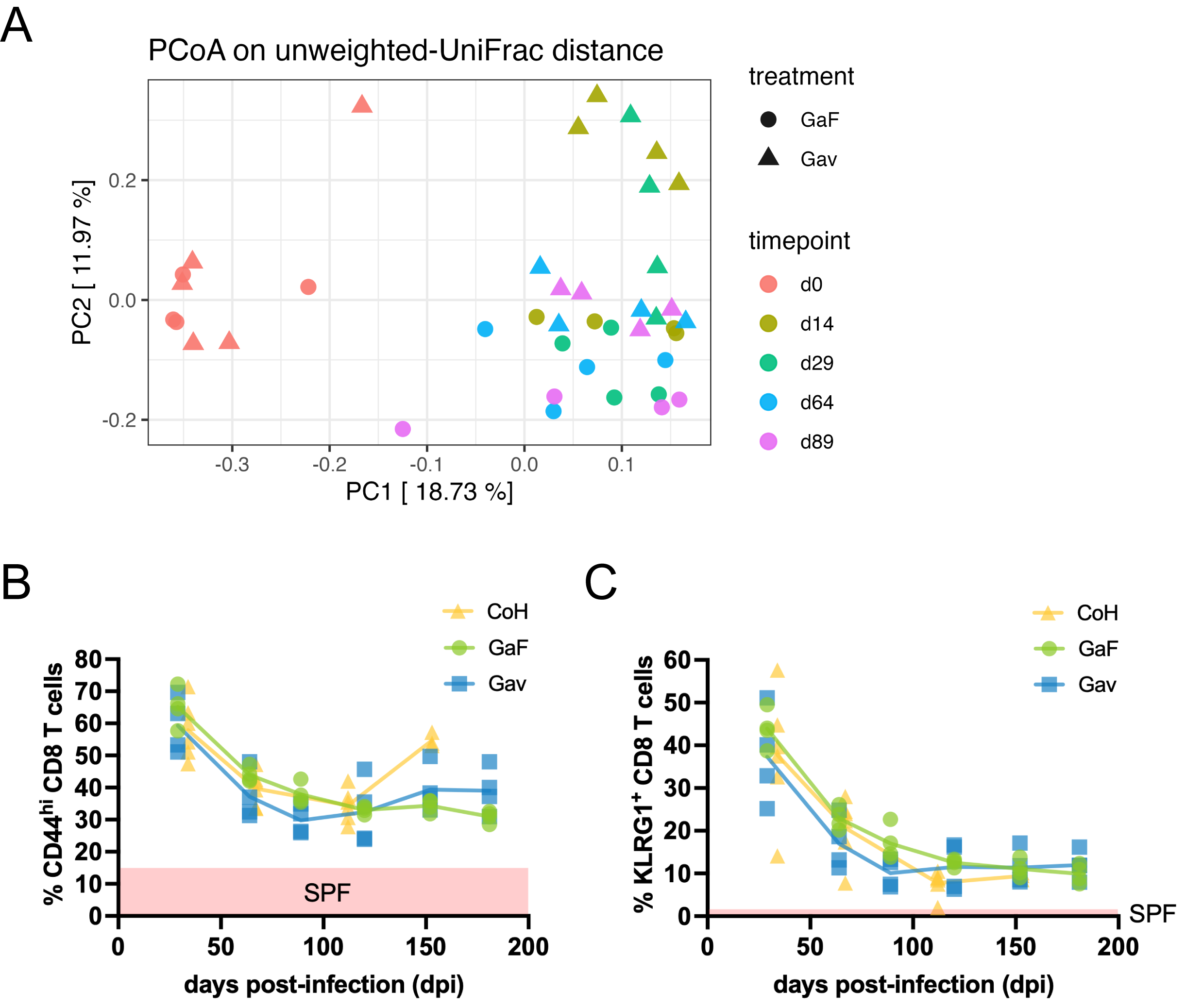

**Figure S1. Long-term stability of gavage/fomite and gavage treatment on the gut microbiome and immune phenotypes.** (A) PCoA plot based on weighted UniFrac distance. Each color represents different time points and each treatment is distinguished by shape. Longitudinal stability of CD44^hi^ CD8^+^ T cells (B) and KLRG1^+^ CD8^+^ T cells (C). The red box indicates the average frequencies of those cell populations in SPF mice. SPF: specific pathogen-free mice, CoH: cohoused laboratory mice with pet store mice, GaF: gavage/fomite-treated mice, Gav: gavage-only mice.

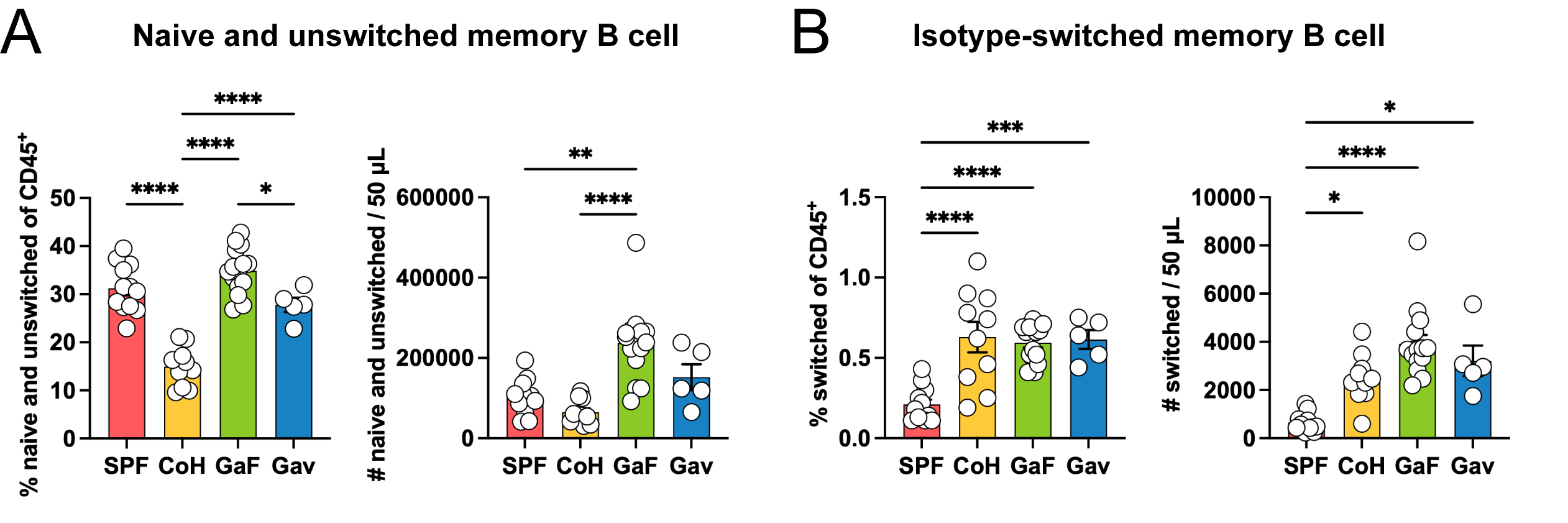

**Figure S2. The B cell subsets of different natural microbiota-based mouse models in the blood.**  Peripheral blood B cell subsets were characterized by flow cytometry ≥ 60 days post-treatment. Cells were gated on CD45^+^ B220^+^ CD38^+^ GL7^-^ populations. Enumeration of naïve and memory B cells (A) and isotype-switched memory B cells (B) frequencies and counts. Each B cell subset was gated as follows: (A) naïve and unswitched memory B cells (IgM^+^ and/or IgD^+^), (B) isotype-switched memory B cells (IgM^-^ IgD^-^). Bars indicate mean ± SEM. Significances were determined by the One-way ANOVA with Tukey’s multiple comparisons test or the Kruskal-Wallis test with Dunn's multiple comparisons test based on the normality of data. *p < 0.05, **p < 0.01, ***p < 0.001, and ****p < 0.0001.

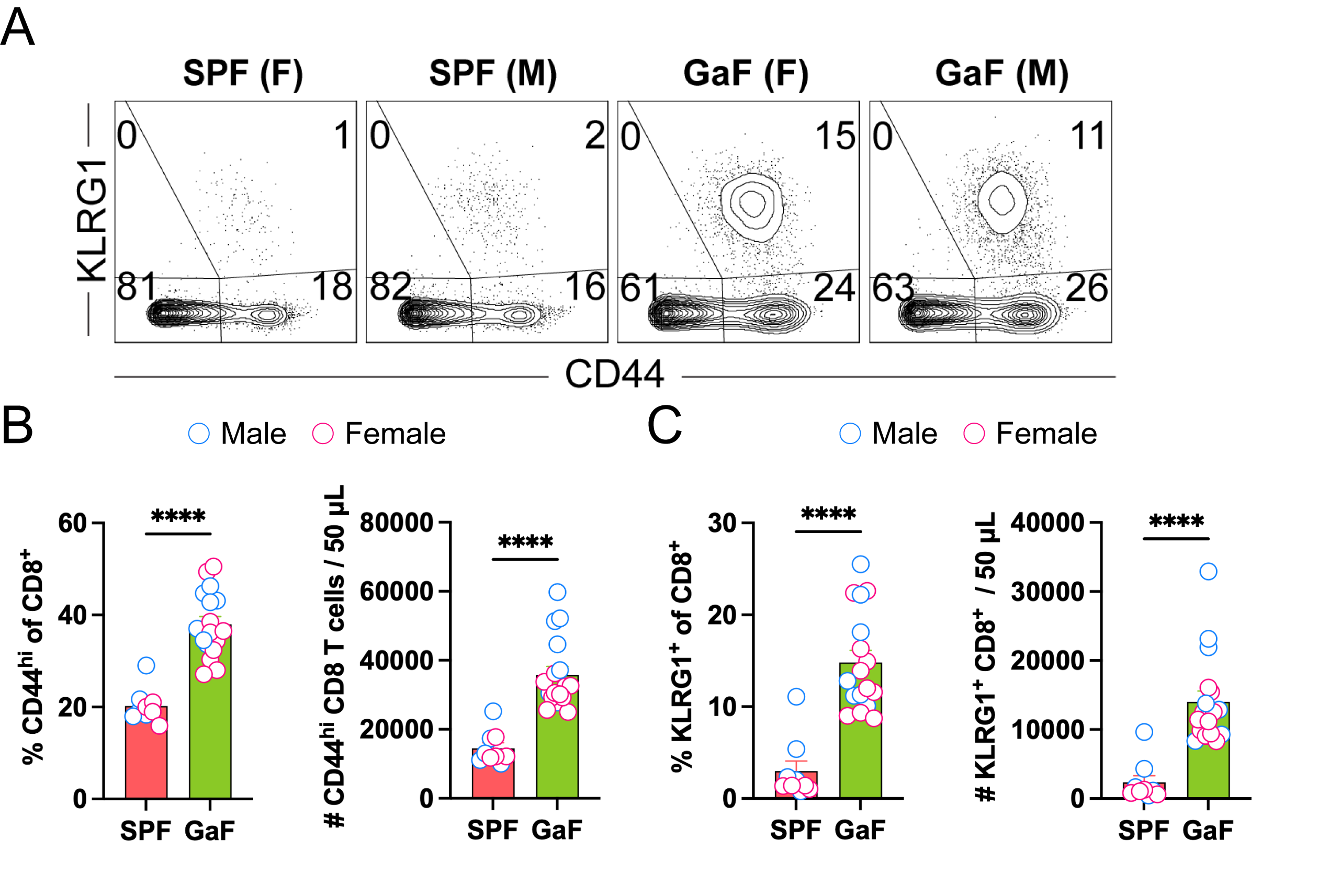

**Figure S3. The effects of gavage/fomite on CD8 T cell differentiation is indistinguishable between male and female mice.** Flow cytometry was used to characterize the peripheral blood immune cells of male gavage/fomite mice. (A) Representative flow plots of CD8^+^ T cells. Cells were gated on CD45^+^ CD11b^-^ B220^-^ CD4^-^ CD8^+^. Enumeration of CD44^hi^ CD8^+^ T cells (B) and KLRG1^+^ CD8^+^ T cells (C) frequencies and counts. All mice were bled at ≥ 60 days after treatment. Circles indicate male (blue) and female (red) mice. Bars indicate mean ± SEM. Significances were determined by the Mann-Whitney test. ****p < 0.0001.

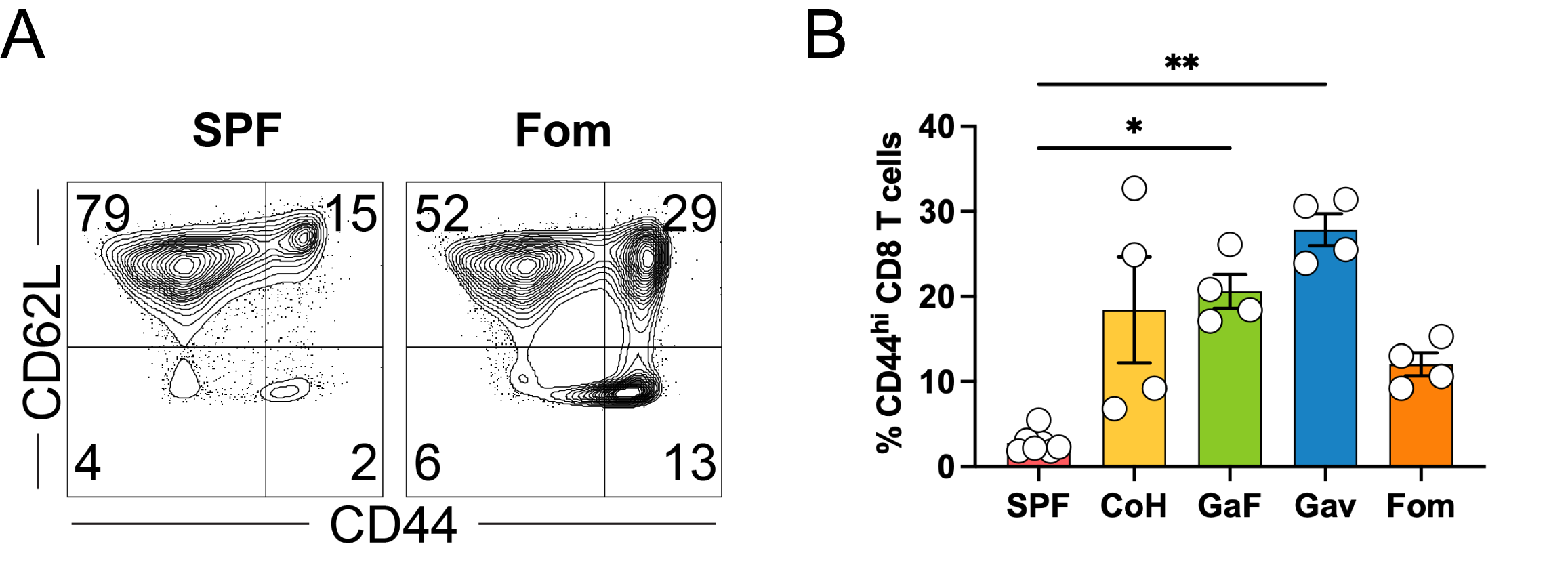

**Figure S4. The effects of fomite-only (Fom) treatment on the CD44^hi^ CD8^+^ T cells in the spleen.** Flow cytometry was used to characterize immune cells in the spleen. (A) Representative flow plots of CD8^+^ T cells. (B) The frequency of CD44^hi^ CD8^+^ T cells in different natural microbiota-based mouse models. Spleens were harvested ≥ 60 days post-treatment. The immune cells were gated as described in Fig. S4. Bars indicate mean ± SEM. Significances were determined by the Kruskal-Wallis test with Dunn's multiple comparisons test.  *p < 0.05 and **p < 0.01.

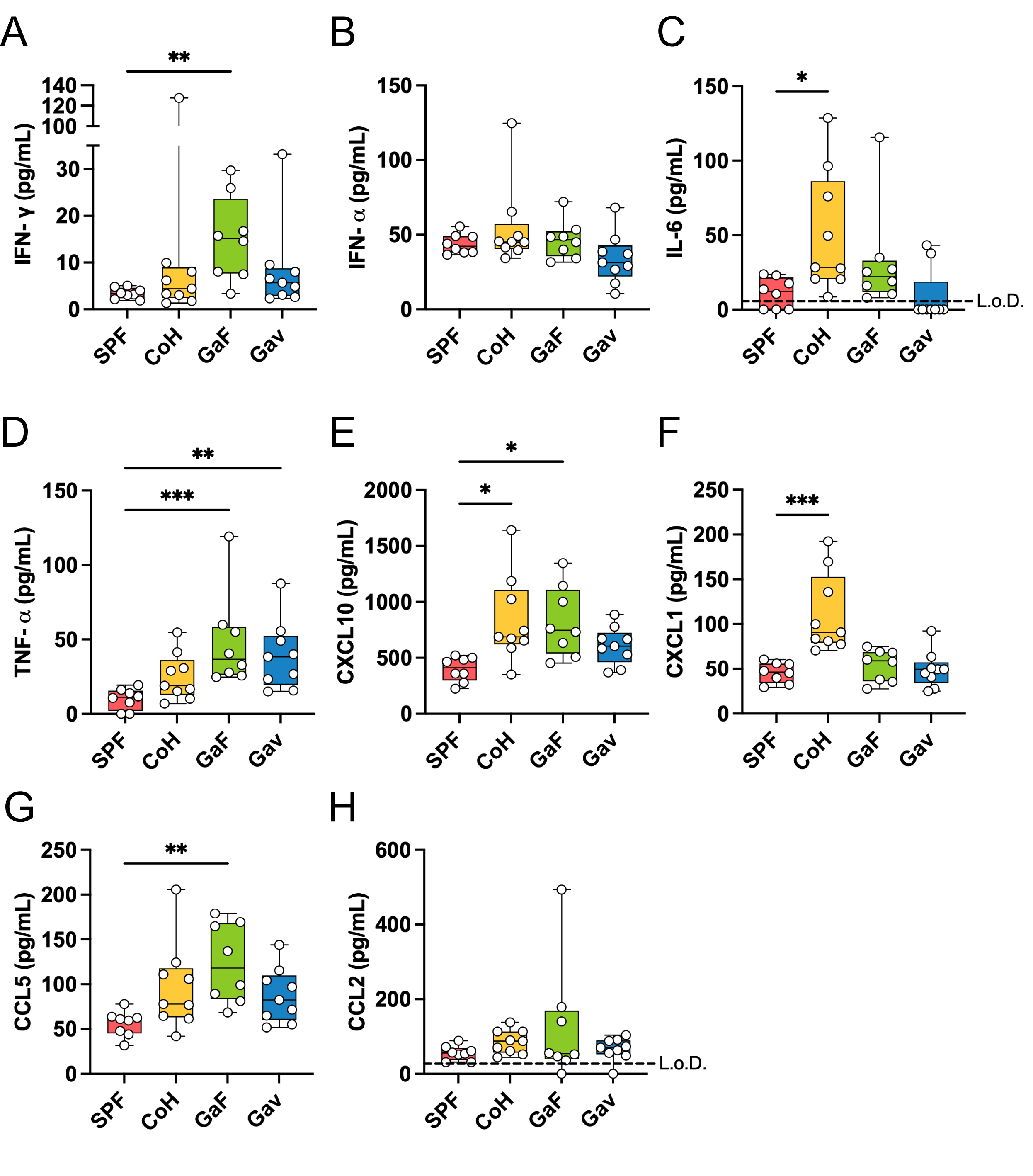

**Figure S5.** **Boxplots of cytokine levels in the serum related to Fig. 3E.** (A) IFN-γ. (B) IFN-α. (C) IL-6. (D) TNF- α. (E) CXCL10. (F) CXCL1. (G) CCL5. (H) CCL2. Significances were determined by the One-way ANOVA with Tukey’s multiple comparisons test or the Kruskal-Wallis test with Dunn's multiple comparisons test based on the normality of data.  *p < 0.05, **p < 0.01, and ***p < 0.001.

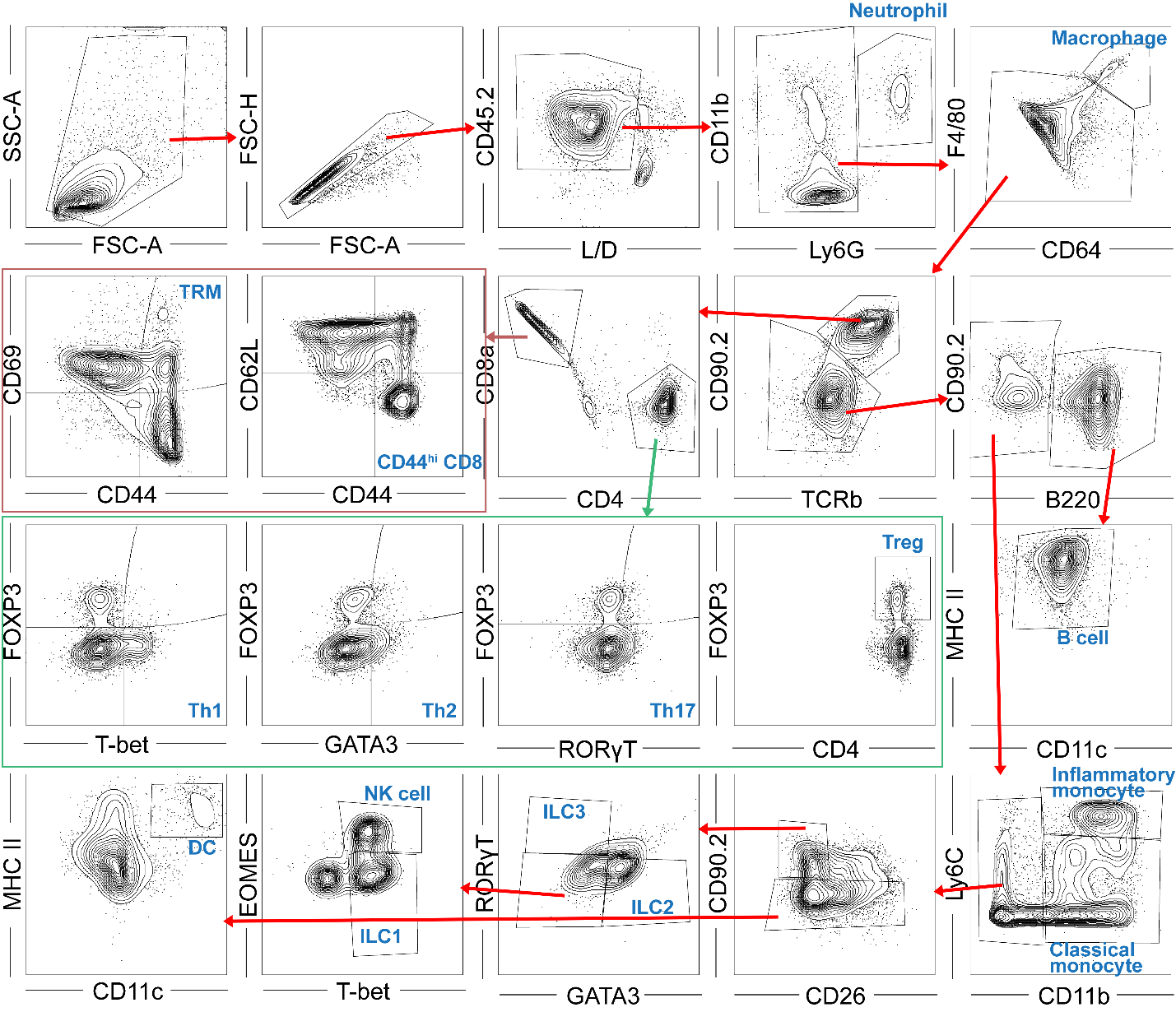

**Figure S6. Gating strategy for t-SNE analysis.**

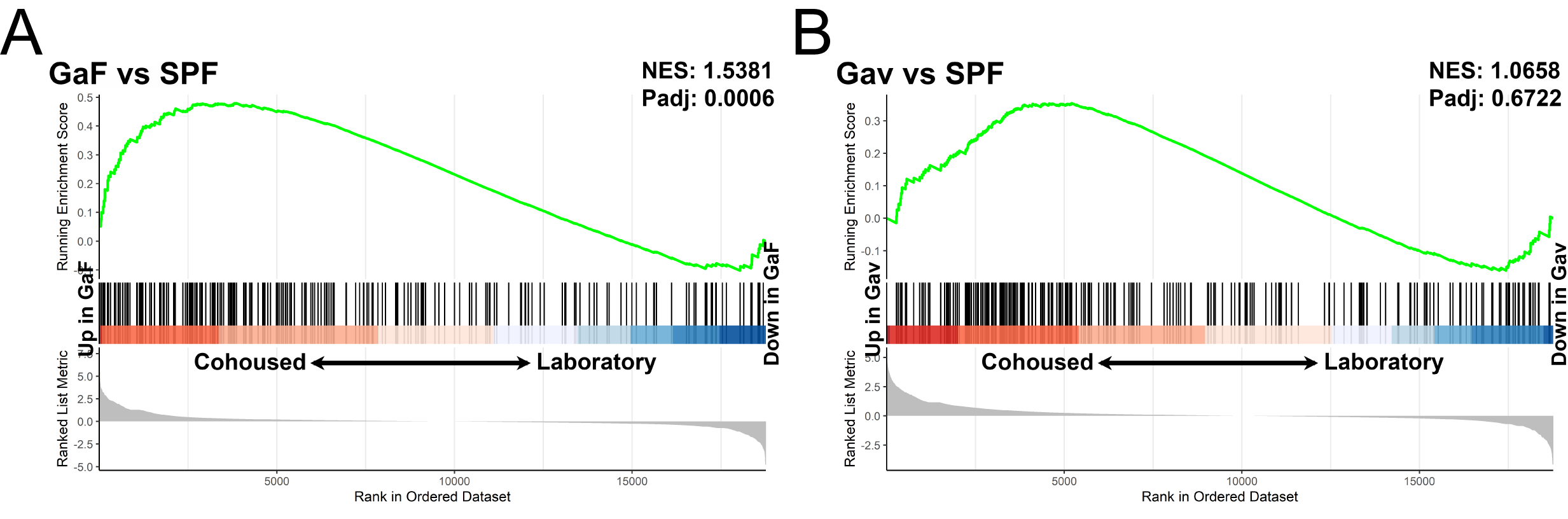

**Figure S7. GSEA plots of GaF vs SPF and gavage-only vs SPF.** Bulk RNA sequencing was performed to compare transcriptional profiles of GaF and gavage-only mice in the spleen to previously published cohoused and laboratory mice PBMC datasets. GSEA plots of GaF vs SPF (A) and gavage-only (Gav) vs SPF (B). Signatures comprise the top 400 genes significantly differentially expressed in cohoused and laboratory mice PBMC (data obtained from GSE78979).
